# Multi-omics integrated analysis reveals a specific phenotype of CD8+ T cell may contribute to immunothromosis via Th17 response in severe and critical COVID-19

**DOI:** 10.1101/2022.07.23.501235

**Authors:** Wen-Xing Li, San-Qi An, Shao-Xing Dai, Zhao-Ming Zhou, Xin Zeng, Guan-Hua Deng, Ying-Ying Huang, Ling-Yu Shen, An-Qi Xu, Yao Lin, Jun-Jun Jiang, Mei-Juan Zhou, Wu Wei, Hao Liang, Dao-Gang Guan, Cheng Zhou

**Affiliations:** Department of Biochemistry and Molecular Biology, Guangdong Provincial Key Laboratory of Single Cell Technology and Application, School of Basic Medical Sciences, Southern Medical University, Guangzhou 510515, Guangdong, China; Biosafety Level-3 Laboratory, Guangxi Collaborative Innovation Center for Biomedicine, Guangxi Medical University, Nanning 530021, Guangxi, China; State Key Laboratory of Primate Biomedical Research, Institute of Primate Translational Medicine, Kunming University of Science and Technology, Kunming, Yunnan 650500, China; Department of Radiation Medicine, Guangdong Provincial Key Laboratory of Tropical Disease Research, School of Public Health, Southern Medical University, Guangzhou 510515, China; Department of Oncology, Guangdong Sanjiu Brain Hospital, Guangzhou 510515, China; CAS Key Laboratory of Computational Biology, Shanghai Institute of Nutrition and Health, University of Chinese Academy of Sciences, Chinese Academy of Sciences, Shanghai 200031, China; Department of Nephrology, Zhujiang Hospital, Southern Medical University, Guangzhou 510280, China; Department of Neurosurgery, Zhujiang Hospital, Southern Medical University, Guangzhou 510282, China; Department of Radiation Oncology, Nanfang Hospital, Southern Medical University, Guangzhou 510515, China

**Author notes:** Corresponding author: Hao Liang, Dao-Gang Guan, Cheng Zhou. These authors have contributed equally to this work (Wen-Xing Li, San-Qi An, Shao-Xing Dai).

**Keywords:** COVID-19, T lymphocyte reduction, Th17 cell differentiation, immunothrombosis

## Abstract

T lymphocyte reduction and immunosenescence frequently occur in severe and critical coronavirus disease 2019 (COVID-19) patients, which may cause immunothrombosis and numerous sequelae. This study integrated analyzed multi-omics data from healthy donors, pneumonia, COVID-19 patients (mild & moderate, severe, and critical), and convalescences, including clinical, laboratory test, PBMC bulk RNA-seq, PBMC scRNA-seq and TCR-seq, BAL scRNA-seq, and lung proteome. We revealed that there are certain associations among T lymphocyte reduction, CD8+ T cell senescence, Th17 immune activation, and immunothrombosis. A specific phenotype (S. P.) CD8+ T cells were identified in severe and critical COVID-19 patients in both PBMC and BAL scRNA-seq, which showed highly TCR homology with terminal effector CD8+ T cells and senescent CD8+ T cells. Pseudotime analysis showed that the S. P. CD8+ T cells were located in the transition trajectory from mild to severe disease. Which may be activated by terminal effector CD8+ T cells or senescent CD8+ T cells, thereby promoting Th17 cell differentiation. This phenomenon was absent in healthy donors, mild and moderate COVID-19 patients, or convalescences. Our findings are an important reference for avoiding the conversion of patients with mild to severe diseases and provide insight into the future prevention and control of COVID-19 and its variants.

## 1. Introduction

Since the coronavirus disease 2019 (COVID-19) outbreak, the understanding of this new respiratory disease has gone through a process from unknown to gradual understanding. During this period, from epidemiological, virological to clinical and case, autopsy, and immunological research, the understanding of the disease has been deepened [1-3]. Difficulty in breathing can occur in the early stage of new coronary pneumonia. With the aggravation of lung infection, patients can experience respiratory failure and even increase the risk of death. A large number of studies have shown that the reduction and exhaustion of cytotoxic lymphocytes (such as CTL cells and NK cells) are important markers of critical illness in COVID-19 and its variants, and the degree of reduction is positively correlated with critical illness and mortality [4, 5]. At the same time, it is worth noting that patients with new coronary pneumonia are more often accompanied by venous thrombosis and abnormal coagulation, and often indicate a poor prognosis [6]. In critically ill patients with COVID-19, the incidence of venous thromboembolism (VTE) and arterial thrombosis ranges from 15% to 30% [7-9]; up to 80% of patients with COVID-19 have small alveoli during ultrasound-guided minimally invasive autopsy. There is a fibrin thrombus in the artery [10]. Autopsy evaluations confirm the widespread presence of pulmonary embolism and multiple microthrombosis in multiple organs, including but not limited to lung, kidney, liver, heart, and brain [11-13]. As an evolutionarily conserved host immune defense mechanism, the role of immune thrombosis and even disseminated intravascular coagulation (DIC) in the pathogenesis of critical patients have been paid more and more attention [14, 15]. However, whether there is an intrinsic relationship between immune failure and abnormal coagulation function in COVID-19 patients is unclear, and the underlying molecular biological mechanism is still lacking in in-depth research.

The results of PBMC single-cell sequencing showed that during the progression of COVID-19, accompanied by a decrease in the number of lymphocytes, an increase in the number of monocytes, and the exhaustion of T cells, a large number of inflammatory cytokines were highly up-regulated in patients with severe COVID-19 [16]. By sequencing the single-cell transcriptomes of different tissues of 196 people (including healthy people, mild COVID-19, severe COVID-19, mild recovery and severe recovery), the researchers found that megakaryocytes and monocytes in PBMC are cytokines in severe patients. A key source of storms and revealed an interaction between hyperinflammatory cell subtypes in lung and PBMC [17]. Analysis of single-cell transcriptome sequencing data of bronchoalveolar lavage fluid showed that the number of T cells, B cells, and NK cells decreased, and the number of macrophages and neutrophils increased in patients with severe COVID-19 compared with patients with mild disease [18]. T cells such as Th17 cells and Th1 cells are highly dysregulated in severe COVID-19 patients, suggesting that Th17 cells play a key role in the progression from mild to severe COVID-19 [19]. In addition, Luminex detection of cytokines in patients with new crowns showed that multiple Th17 cell-related cytokines (IL-1α, IL-1β, IL-6, IL-17, THFα, IFNγ, etc.) Significantly high profile [20]. Th17 immune response is an important factor in activating centriocytes and triggering NETosis, which leads to immune thrombosis in multiple organs (including lung, liver, heart, kidney, etc.) [21]. Therefore, inhibiting Th17 cell activation and reducing NETosis and immune thrombosis may become a new strategy to reduce the transformation rate of critical illness and reduce the large-scale medical burden.

This study integrated multi-omics data including clinical and experiment data, bulk RNA-seq, scRNA-seq and proteomic data, which contains COVID-19 patients with mild and moderate, severe and critical disease, and patients in the convalescent stage, as well as healthy controls and pneumonia. We reveal that decreased T lymphocytes were negatively correlated with increased D-dimer in COVID-19 patients, this may lead to an increase in cellular senescence in severe and critical COVID-19 patients. We identified a specific phenotype (S. P.) CD8+ T cells only in severe and critical stage, the S. P. CD8+ T cells may be affected by the senescent cells and promote Th17 cell differentiation, thereby triggering a series of immune responses and cause immunothrombosis. This study has important clinical and public health value in inhibiting coagulation disorders and reducing the conversion rate of critical patients, thereby avoiding large-scale medical burdens.

## 2. Methods

### 2.1 Transcriptome and clinical data of COVID-19

Clinical information, laboratory test and transcriptome data of COVID-19 patients were acquired from the previous report [22]. Which contains blood routine test data of 1551 patients, plasma cytokine and complement data of 48 patients in mild & moderate, severe, and critical COVID-19, as well as PBMC bulk RNA-seq for healthy, pneumonia, and COVID-19 patients in different stages.

### 2.2 Bulk RNA-seq data processing

Sequenced RNA-seq data (paired-end reads) of “fastq” format were processed with Skewer (version: 0.2.2) [23] for quality control and adapter trimming. Then used Pairfq (version: 0.17.0, https://github.com/sestaton/Pairfq) for reads pairing and removing the unpaired reads in each filtered paired-end file. The generated clean data with high quality were used for further analysis. Human reference genome and gene model annotation files (genome assembly: GRCh38.90) were downloaded from Ensemble database (http://asia.ensembl.org/index.html). The genome index was build using the python scripts included in the HISAT2 package (version: 2.1.0) [24, 25]. The paired-end clean reads were aligned to the reference genome using HISAT2 and run with the default parameters. The aligned file in “sam” format were sorted and converted to “bam” format by using SAMtools (version 1.6) [26]. Transcripts assembly and estimates expression levels of the sorted reads were using StringTie (version 1.3.3b) [27] and run with the default settings. Next, the FPKM (Fragments Per Kilobase of transcript per Million fragments mapped) of each gene was calculated and summarized in “Ballgown” package (version 2.2.0) [28].

### 2.3 Gene annotation and filtration

Human gene annotation file was obtained using BioMart tool in Ensemble database [29]. Gene annotation of processed RNA-seq file were using custom written R scripts. There were 44 types of RNAs in the annotated expression matrix and we extracted mRNAs (type of “protein coding”) for further analysis. To ensure the data quality, A gene with a log2 FPKM expression greater than 0.1 in more than 90% samples was retained. Finally, we got an expression matrix with 12328 unique genes and 25 samples.

### 2.4 Differential expression analysis

Differentially expressed gene analysis was performed using the empirical Bayes algorithm in the “limma” package [30] in R statistical software version 4.0.3. The differential expression fold changes (FC) were represented by the log2 transformation. The differentially expressed genes (DEGs) were determined between different disease severity groups with the criteria that the false discovery rate (FDR) corrected P value < 0.05 and |log2FC| > 1.0. Hierarchical cluster analysis was performed on the DEGs among different groups utilizing the “ward.D2” algorithm in the “pheatmap” package.

### 2.5 Clinical data analysis

The results for the categorical variables (i.e., gender) are presented as numbers and percentages of cases. The continuous variables (i.e., age, blood routine, blood chemistry, blood gas analysis, and blood coagulation) are presented as the median (interquartile range). Pearson correlation analysis is used to compare the correlation between two clinical features. The comparison of continuous variables in two groups used the independent samples t-test. The comparison of continuous variables in three or more groups used the one-way ANOVA with LSD Post Hoc test. A P value < 0.05 was considered significant.

### 2.6 Cell type abundance analysis

Cibersort web server (https://cibersort.stanford.edu/) [31] was used to identify different cell types and their relative abundance. The signature gene file was chosen as LM22 reference sample file. We manually merged similar cell types in LM22, and generated a dataset contains 12 cell types: B cells, dendritic cells, eosinophils, macrophages, mast cells, monocytes, neutrophils, NK cells, plasma cells, T cells CD4, T cells CD8, and T cells cytotoxic. The average gene expression signatures of the original LM22 cell types were used as the signatures of the corresponding merged cell types.

### 2.7 Trend analysis of immune cell related gene signatures

Expression trend analysis of immune cell related genes were using Short Time-series Expression Miner (STEM, version: 1.3.13) [32]. The mean expression value of Cibersort reference genes in healthy, COVID-19 mild & moderate, COVID-19 severe, and COVID-19 critical were used as input parameters. Other parameters use default settings. The software generates multiple model profiles based on gene expression features. Each profile ordered based on the P-value significance of number of genes assigned versus expected. A P value < 0.05 was considered significant.

### 2.8 KEGG and GO enrichment analysis

The reference human genes and pathways were obtained from the Kyoto Encyclopedia of Genes and Genomes (KEGG) database (http://www.kegg.jp/) [33]. The information on human genes and related GO biological functions were downloaded from the QuickGO database (http://www.ebi.ac.uk/QuickGO-Beta/) [34]. Pathways with less than 10 genes were removed. The enrichment analysis was performed using the hypergeometric test. An FDR-corrected P value < 0.05 was considered significantly enriched. In this study, we mainly focus on pathways related to the immune system and signal transduction.

### 2.9 Screening of hub genes

Hub genes were screened from immune system and signal transduction pathways. We firstly calculated the mean expression of gene in each group (healthy, pneumonia, mild & moderate, severe, and critical COVID-19). Then the ratio of each gene was calculated using the following formula:

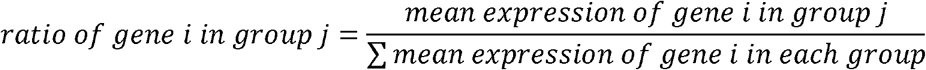

Next, the top 10 genes according to the descending order of the ratio in each group were screened and defined as the hub genes.

### 2.10 Construction of transcriptional regulatory networks

Human transcription factors (TFs) and corresponding transcriptional regulatory relationship were downloaded from the TRRUST webserver v2.0 [35]. Pearson’s correlation analysis was used to calculate the correlation between TFs and target genes. TF-gene pairs with correlation coefficient high than 0.8 and P-value less than 0.05 was considered to be significantly correlated and were used to constructed the TF-gene regulatory networks (TRNs). Network visualization was used Cytoscape software (version 3.9.2) [36] and the NetworkAnalyzer tool was used to analyze the TRNs.

### 2.11 PBMC and BAL scRNA-seq data analysis

Dataset of GSE158055 was downloaded from the NCBI GEO database, which contains 1.46 million cells from 196 COVID-19 patients and healthy donors [17]. Cells derived from peripheral blood mononuclear cells (PBMC) samples (including fresh PBMC and frozen PBMC) were used for further analysis. The standard pre-processing workflow for scRNA-seq data in Seurat [37] was used for quality control, data normalization, feature selection, data scaling, linear dimensional reduction, and t-SNE clustering of the cells in PBMC samples. Cell annotation was using the SingleR (version 1.6.1) and celldex (version 1.2.0) [38]. The reference dataset was chosen the Monaco immune data [39], which contains 8 types of CD4+ T cells including T regulatory cells (Treg), Th1 cells, Th1/Th17 cells, Th17 cells, Th2 cells, follicular helper T cells (Tfh), naive CD4+ T cells, and terminal effector CD4+ T cells (TEF CD4+), and 4 types of CD8+ T cells including naive CD8+ T cells, central memory CD8+ T cells (TCM CD8+), effector memory CD8+ T cells (TEM CD8+), and terminal effector CD8+ T cells (TEF CD8+). Bronchoalveolar lavage (BAL) scRNA-seq data of 5 patients with mild and 26 with critical COVID-19 patients was downloaded from the European Genome-Phenome Archive database (EGAS00001004717) [19] and analyzed follow the above steps.

### 2.12 Lung proteome data analysis

Lung proteomic data of healthy and COVID-19 patients were collected from the iProX database (https://www.iprox.cn/) with accession ID: IPX0002393000 [21]. The lung protein matrix of healthy and COVID-19 patients was used for further analysis. We acquired proteins in the following pathways in the Enrichr database [40]: thrombosis (DisGeNET), blood coagulation (GO: 0007596), lung inflammation (MP: 0001861), lung fibrosis (WP3624), neutrophil activation (GO: 0042119), Th1 and Th2 cell differentiation (hsa04658), Th17 cell differentiation (hsa04659), and T cell activation (GO: 0042110).

### 2.13 Cell senescence status analysis

Human senescence-related genes were downloaded from the Human Ageing Genomic Resources database (https://genomics.senescence.info/). In order to evaluate the senescence status of CD8+ T cells, cellular senescence status was calculated as the mean expression of senescence-related genes in each cell. All CD8+ T cells were sorted in ascending order according to the cellular senescence status, and the top 10% of cells were defined as the senescent CD8+ T cells.

### 2.14 Pseudotime analysis

Pseudotime analysis was performed using the R package Monocle 3 [41]. The single-cell transcriptome data of PBMCs of COVID-19 was used to construct the pseudo-time trajectories of total T cells in healthy controls, COVID-19 patients, and convalescences. For each analysis, PCA-based dimension reduction was performed with differentially expressed genes of each phenotype, followed by two-dimensional visualization with UMAP. We retained the main trajectories and removed the fragmented branch in each analysis. The healthy controls or T cells in the initial phase of the immune response were defined as a root state for the calculation of the immune response process and pseudotime.

### 2.15 TCR homology analysis

T cell receptor (TCR) homology analysis was performed using the TCR data from COVID-19 patients corresponding to single-cell transcriptome samples [17]. The TRUST4 algorithm [42] was able to reconstruct the immune receptor repertoire in T and B cells from RNA-sequencing data. We used the TRUST4 package to calculate the homology scores of T cell subtypes. Cells with a TCR homology score higher than 80 are considered highly homologous.

### 2.16 Cell-cell communication analysis

Cell-cell communication analysis of different cell subtypes were using the CellChat software package [43]. CellChat is able to compare the number and strength of interactions between different cell populations, define cell-specific signaling networks and calculate the afferent and efferent signals associated with each cell population, and screen for crucial ligand/receptors in different cell types.

## 3. Results

### 3.1 Immune cell abundance and transcriptome profiles in COVID-19 severe and critical patients

This study integrated analyzed multi-omics data from healthy donors, pneumonia, COVID-19 patients (mild & moderate, severe, and critical), and convalescences, including clinical, laboratory test and PBMC bulk RNA-seq [22], PBMC scRNA-seq and TCR-seq [17], BAL scRNA-seq [19], and lung proteome [21] (Figure 1A). Clinical data showed that lymphocytes and platelets decreased significantly as the COVID-19 progressed and were much lower than in healthy donors, whereas the levels of neutrophil and white blood cell (WBC) showed the opposite trend (Figure 1B, Table S1). Other clinical indicators, such as D-dimer was dramatically increased with disease progression and significantly higher than in healthy donors (Table S1). Curve fitting revealed a decreased T lymphocyte was significantly negatively correlated with D-dimer in COVID-19 patients (Figure 1C). We used Cibersort [31] to estimate the abundance of immune cells based on PBMC bulk RNA-seq data. The relative abundances of neutrophils and macrophages were mainly distributed in severe and critical COVID-19 patients, whereas CD4+ T cells, B cells, and monocytes were mainly distributed in mild & moderate COVID-19 patients (Figure 1D). Especially, the relative abundance of neutrophils gradually increased whereas monocytes gradually decreased with disease progression. CD4+ T cells and CD8+ T cells were significantly decreased in critical compared with mild & moderate in COVID-19 patients (Figure 1E). Gene clusters with different expression trends of the above immune cell-related genes were obtained by the Short Time-series Expression Miner analysis [32]. The expression trend and enriched GO biological functions of gene cluster 1 and 2 was consistent with the abundance change of neutrophils and CD4+ T cells, respectively (Figure 1F). Combining transcriptomic and clinical performance, there were serious decreased T lymphocyte, increased neutrophils, and coagulation abnormalities in severe and critical COVID-19 patients.

**Figure 1.**
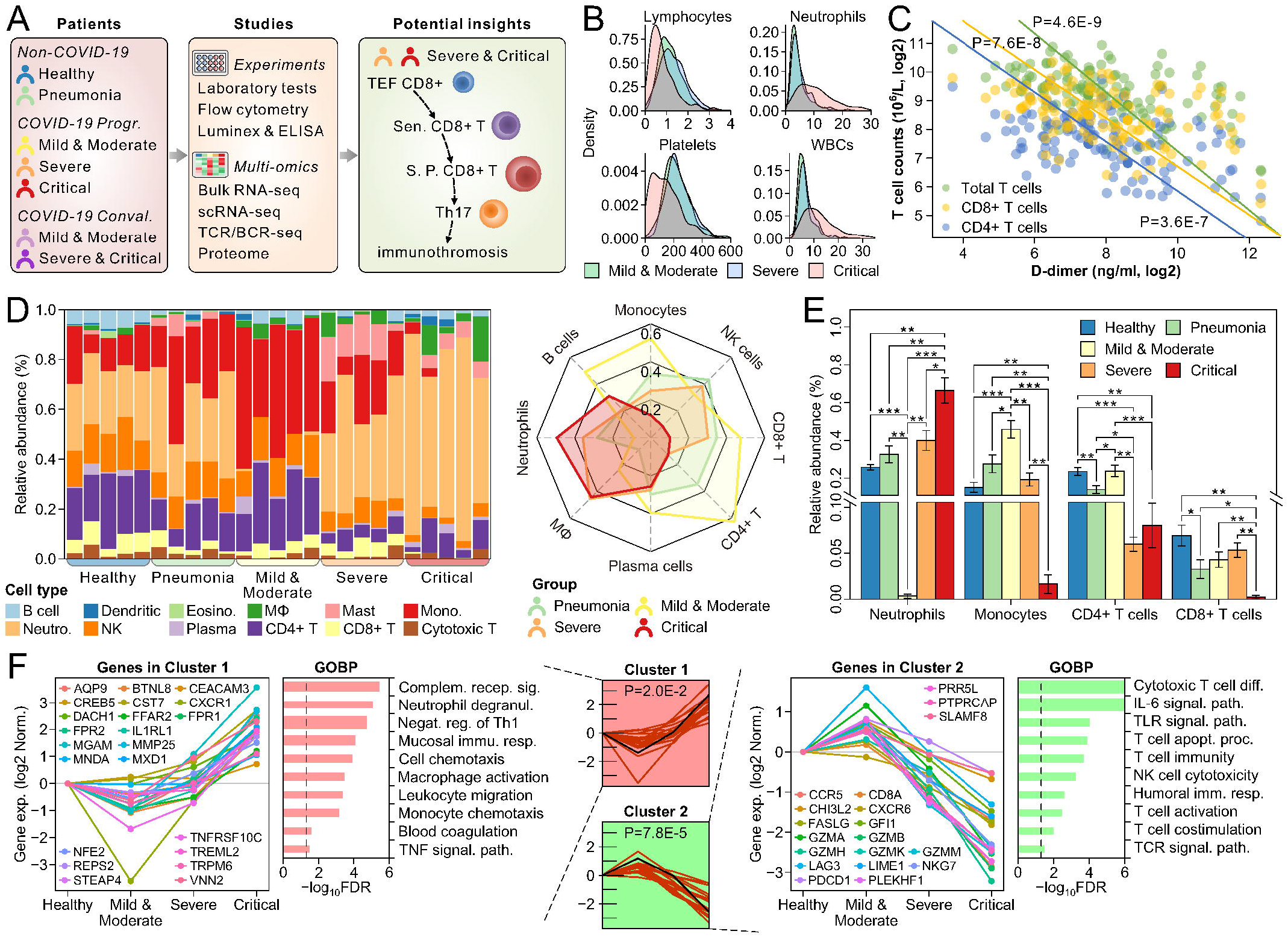
Immune cell abundance and transcriptome features in COVID-19 patients. (A) Flowchart describing the experimental design of this study. (B) The distribution of lymphocytes, neutrophils, platelets, and white blood cells (WBCs) in mild & moderate, severe, and critical COVID-19 patients. (C) Correlation of T cell counts and plasma D-dimer levels. (D) Relative abundance of 12 types of immune cells in each sample (left) and the normalized average abundance in four groups (right). (E) ANOVA and LSD post hoc tests of cellular abundance of neutrophils, monocytes, CD4+ T cells, and CD8+ T cells in healthy control, pneumonia, and COVID-19 patients. (F) Expression trends and enriched GO biological functions (BP) of the representative two gene clusters. Significance: * P < 0.05, ** P < 0.01, *** P < 0.001.

### 3.2 Th17 cells and coagulation function play important roles in COVID-19 progression

We calculate the differentially expressed genes (DEGs) and performed KEGG pathway enrichment analysis in the pairwise comparison among healthy donors, pneumonia, mild & moderate, severe, and critical COVID-19 patients. Multiple immune system and signal transduction related pathways were significantly enriched in critical COVID-19 patients compared with healthy donors, pneumonia, and severe COVID-19 patients (Figure 2A, 2B). We screened hub genes among these five groups and constructed the hub gene-pathway network (Figure 2C). The results showed that Th17 cell differentiation, IL-17 signaling pathway, and complement and the coagulation cascade pathways were significantly enriched. Among them, RORC is a key transcription factor driving Th17 cell differentiation, which is highly expressed in mild & moderate COVID-19; IL1R1 is over-expressed in critical COVID-19. FOSB, a key gene of the IL17 signaling pathway, was dramatically up-regulated in mild & moderate COVID-19 compared with other groups; MMP9 was highly expressed in severe and critical COVID-19. Consistent with the results of transcriptome analysis, Luminex and ELISA results showed that multiple cytokines (IL-17A, IL-21, IL-22, IL-23, IL-6, IL-1α, IL-1β, IL-1RA, GM-CSF, IFNγ, IL-12p70, and TNFα) and complements (C4a, C4b, C5a, and C5b) were gradually upregulated with disease progression (Figure 2D). These findings herald a strong association between Th17 immune response and coagulation abnormalities in severe and critical COVID-19 patients. We further constructed a transcriptional regulatory network of the genes in Th17 cell differentiation, IL-17 signaling pathway, and complement and the coagulation cascade pathways (Figure 3A). Almost all TFs were positively correlated with target genes, and these genes were up-regulated in mild & moderate COVID-19 patients compared with healthy donors and pneumonia. Several crucial TFs (such as FOS, NFKB1, NFKBIA, RARA, STAT3) were high-expressed in critical COVID-19 patients compared with severe patients. Pearson’s correlation analysis showed that the representative genes were positively correlated with monocytes and CD4+ T cells, whereas negatively correlated with neutrophils (Figure 3B). Next, we will explore the interaction and function of different cell types in severe and critical COVID-19 patients.

**Figure 2.**
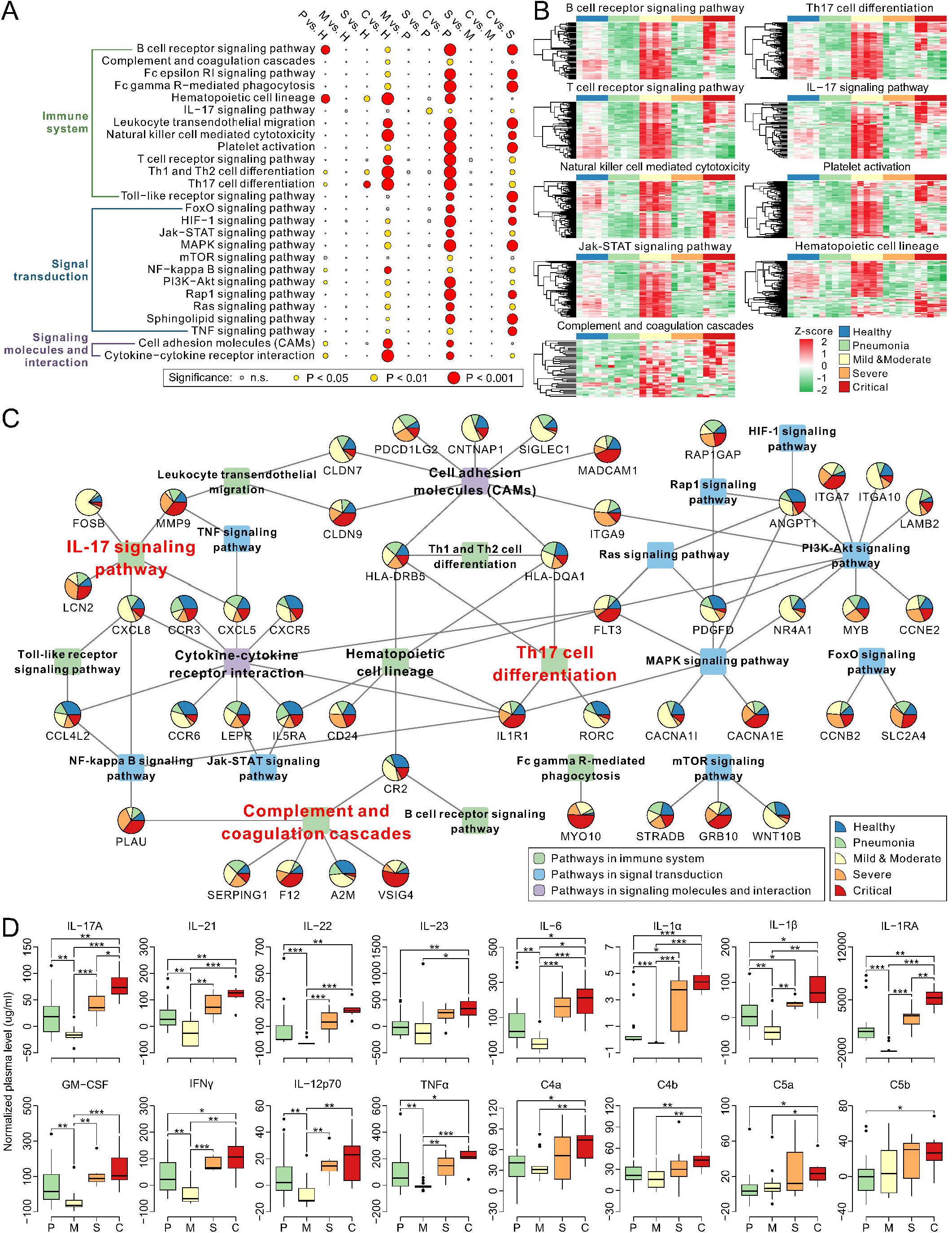
Hub genes and enriched immune-related pathways in COVID-19 patients. (A) Enriched immune system and signal transduction pathways. (B) Heatmap of gene expression profiles in the representative pathways. (C) Network of enriched immune system and signal transduction pathways and hub genes. (D) Difference of cytokines, chemokines, and interleukins in pneumonia, mild & moderate, severe, and critical COVID-19 patients. Significance: * P < 0.05, ** P < 0.01, *** P < 0.001, **** P < 0.0001.

**Figure 3.**
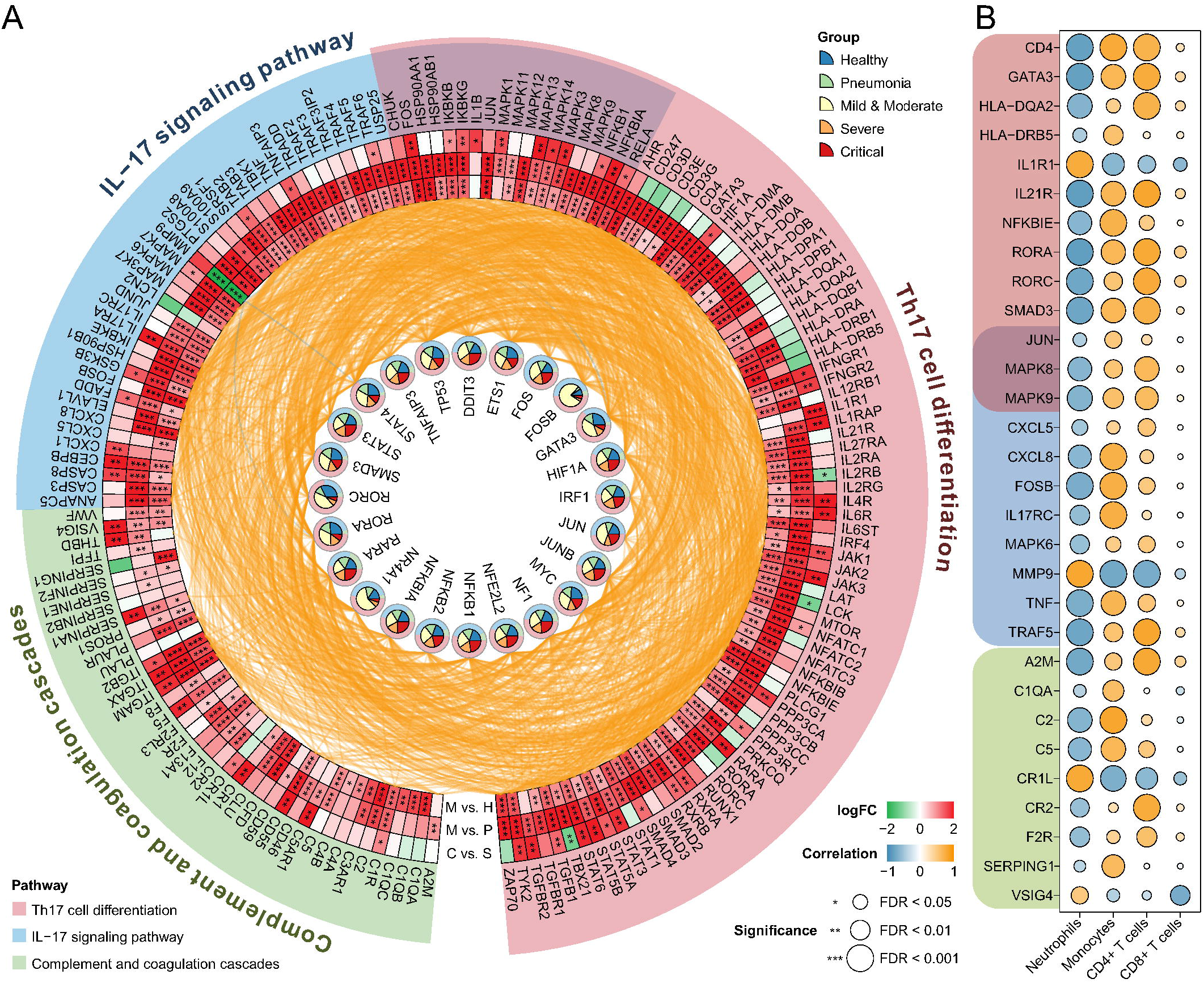
Transcriptional regulatory relationships and gene expression differences in the screened pathways. (A) Correlation of transcription factors and genes in the Th17 cell differentiation, IL−17 signaling pathway, and complement and coagulation cascades pathways. (B) Representative genes significantly correlated with neutrophils, monocytes, CD4+ T cells, or CD8+ T cells. Significance: * P < 0.05, ** P < 0.01, *** P < 0.001.

### 3.3 Expression and functional changes in Th17 cells and senescence CD8+ T cells

We identified 8 subgroups of CD4+ T cells and 4 subgroups of CD4+ T cells (Figure 4A) with PBMC scRNA-seq data of COVID-19 patients [17] by singleR and celldex tools [38]. The results showed that the proportion of Th17 cells in severe and critically COVID-19 patients was significantly higher than that in mild and moderate COVID-19 patients, whereas the proportion of TEF CD8+ cells showed the opposite trend (Figure 4B). Curve fitting revealed a significant negative correlation between Th17 cells and TEF CD8 cells (Figure 4C). We further analyzed the DEGs and enriched pathways of TEF CD8+ cells in COVID-19 progression and convalescence compared with healthy donors, the results showed that the largest number of DEGs, and the cellular senescence, neutrophil extracellular traps formation, IL-17 signaling pathway, Th17 cell differentiation, and T cell receptor signaling pathway were significantly enriched in severe and critical COVID-19 patients (Figure 4C). Human aging genes was downloaded from the Human Ageing Genomic Resources database (https://genomics.senescence.info/), the largest number of senescence DEGs was also found in severe and critical COVID-19 patients (Figure 4C). There were 240 unique DEGs in severe and critical COVID-19 patients compared to healthy donors, multiple senescence genes (CDKN1A, FOS, HIF1A, HSPD1, IL7R, JUN, JUND, NFE2L2, NFKBIA, STAT3, TNF) were highly expressed in severe and critical COVID-19 patients compared to mild and moderate COVID-19 patients and convalescences (Figure 4D). We calculated the senescence status of each CD8+ T cells based on the mean expression of senescence genes, the results showed that the cell senescence state of severe and critical COVID-19 patients was much higher than mild and moderate COVID-19 patients and convalescences (Figure 4D). Therefore, we speculate that there is a certain correlation between the senescence of CD8+ T cells and Th17 cell differentiation. Furthermore, several T cell activation genes were highly expressed in mild and moderate COVID-19 patients, whereas multiple senescence genes and Th17 cell differentiation genes were highly expressed in severe and critical COVID-19 patients. These proteins were also up-regulated in lung proteome of COVID-19 patients compared to healthy (Figure 4E). Based on the constructed transcriptional regulatory relationship, we calculated the mean TF correlation of the genes in pathways of Th17 cell differentiation, IL-17 signaling pathway, and cytokine-cytokine receptor interaction, the above genes were positively regulated by these TFs in severe and critical COVID-19 (Figure 4F). Correlation analysis of multiple gene clusters indicated that multiple genes were shared involved in Th17 cell differentiation, senescence, and blood coagulation (Figure 4G).

**Figure 4.**
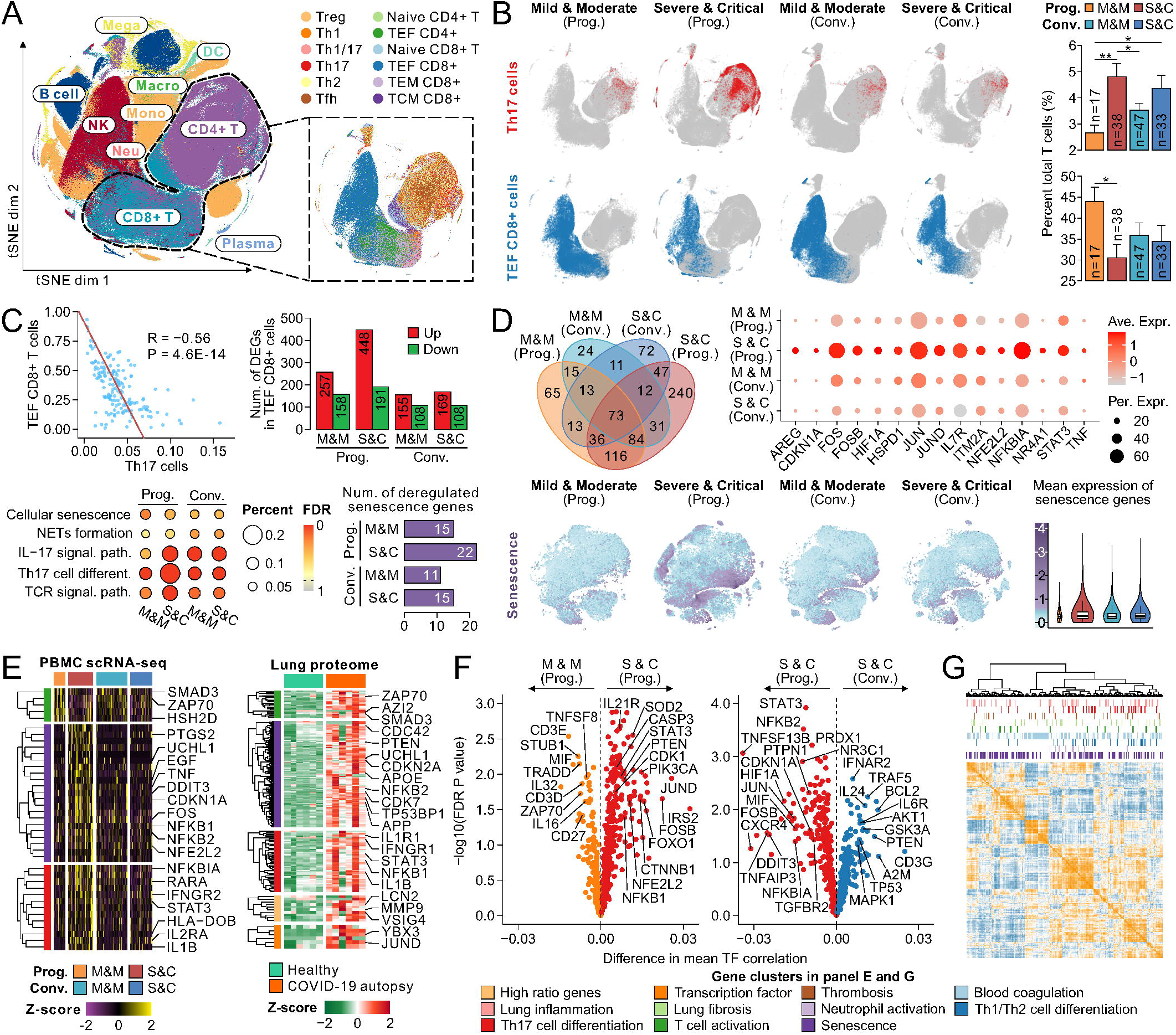
Cell count and gene expression profiles in COVID-19 progression and convalescence stages. (A) Cell clusters in PBMCs of COVID-19 patients and healthy controls. The classification of cell groups in the left panel was according to pervious research [17]. The right panel is the t-SNE cluster of CD4+ and CD8+ T cells. (B) Distribution (left) and differences (right) of Th17 and TEF CD8+ cells in COVID-19 patients and healthy controls. (C) Correlation between Th17 cells and TEF CD8+ cells (top left), differentially expressed genes (DEGs) between COVID-19 patients and healthy controls (top right), enriched pathways (bottom left), and the number of deregulated senescence genes (bottom right) in TEF CD8+ cells. (D) Venn diagram of DEGs (top left) and the expression profiles of representative DEGs (top right) in TEF CD8+ cells, distribution (bottom left) and differences (bottom right) of cellular senescence status in COVID-19 patients and healthy controls. (E) Expression profiles of representative gene clusters in COVID-19 patients and healthy controls. (F) Differences of transcriptional regulatory status in COVID-19 progression and convalescence stages. (G) Pearson correlation of representative gene clusters. Abbreviations: M & M, mild & moderate; S & C, severe and critical; Prog., progression; Conv., convalescence. Significance: * P < 0.05, ** P < 0.01, *** P < 0.001.

### 3.4 The identification of S. P. CD8+ T cells and its roles in Th17 cells functional activation

Pseudo-time analysis of the PBMC scRNA-seq data showed that a subset of cells mainly existed in severe and critical COVID-19 patients, and almost none in healthy donors, mild and moderate COVID-19 patients and convalescence (Figure 5A). Unsupervised clustering showed that a small group of cells in the above subset were highly expressed T cell receptor-ligand genes, whereas other cells showed no this phenotype. TCR homology analysis showed that this group of cells had a high homology with TEF CD8+ and senescence CD8+ T cells. Therefore, we defined the group of cells as the specific phenotype (S. P.) CD8+ T cells (Figure 5B). To explore the function of S. P. CD8+ T cells, pseudo-time analyses were performed on healthy people, mild and moderate, and severe and critical COVID-19 patients. The trajectory showed that S. P. CD8+ T cells were mainly distributed at the junction of mild and moderate and severe and critical COVID-19 patients, and were beside TEF CD8+ cells (Figure 5C). We also identified S. P. CD8+ T cells from BAL scRNA-seq data [19], and the up-regulated genes of S. P. CD8+ T cells in BAL were consistent with those in PBMC (Figure 5D, Table S2). Interestingly, as the number of Sen. CD8+ T cells increased, the number of S. P. CD8+ T cells and Th17 cells also increased gradually only in severe and critical COVID-19 patients (Figure 5E). KEGG pathway analysis of TEF CD8+ cells, Sen. CD8+ T cells, S. P. CD8+ T cells, and Th17 cells showed that Th17 cell differentiation, IL-17 signaling pathway, cellular senescence, platelet activation, and multiple signaling transduction pathways were significantly enriched (Figure 5F). Further pseudo-time analysis of the above four cells in severe and critical COVID-19 patients showed that TEF CD8+ cells were the starting point at the pseudo-time sequence, along the trajectory are Sen. CD8+ T cells, S. P. CD8+ T cells, and Th17 cells (Figure 5G). Cell-cell communication analysis of these different cell types proved that S. P. CD8+ T cells were correlated with TEF CD8+, Sen. CD8+ T cells, and Th17 cells (Figure 5H). Based on the above results, we constructed the specific transcriptional regulatory networks among these four cells (Figure 5I). These results revealed that S. P. CD8+ T cells promote Th17 cell differentiation, and subsequently caused immunothromosis via multiple immunological responses (including complement activation and blood hypercoagulation; platelet activation and fibrin clot formation; neutrophil activation, degranulation and NETosis; and lung injury and vascular inflammation) in severe and critical COVID-19 patients (Figure 6).

**Figure 5.**
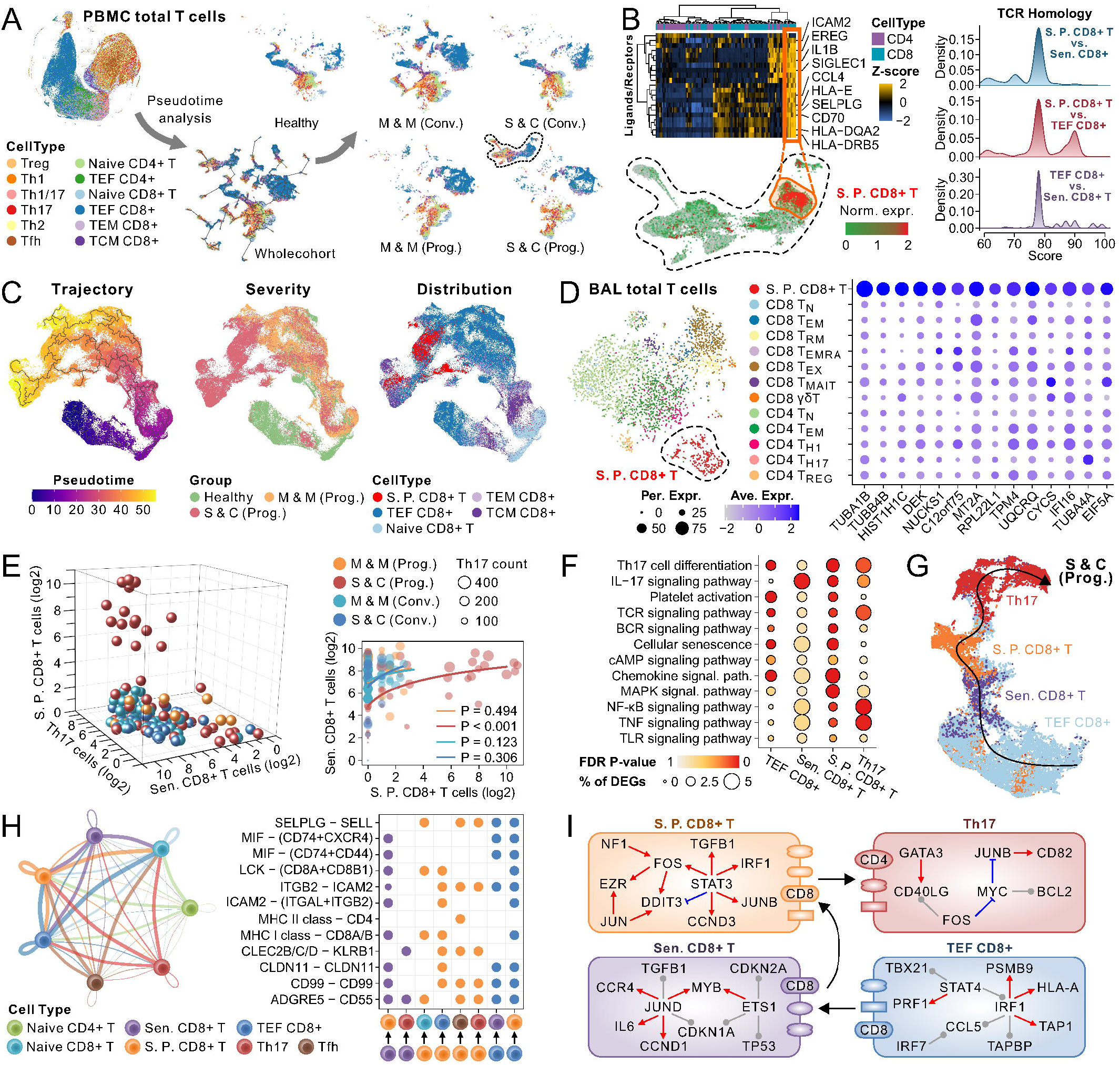
Identification of the specific phenotype (S. P.) CD8+ T cells and its role in severe & critical COVID-19 patients. (A) A unique cell subset was found in PBMC of severe & critical COVID-19 patients. (B) There is a group of cells in the unique cell subset with up-regulated T cell ligand/receptor genes and was defined as S. P. CD8+ T cells (left), and showed high TCR homology with TEF CD8+ cells and Sen. CD8+ T cells (right). (C) S. P. CD8+ T cells is distributed at the stage from mild & moderate to severe & critical of COVID-19 patients. (D) S. P. CD8+ T cells was also found in bronchoalveolar lavage (BAL) of critically COVID-19 patients, and consistent with multiple marker genes of this cell type in PBMC. (E) Cell count correlation of S. P. CD8+ T cells, Th17 cells, and Sen. CD8+ T cells in each sample. There is a significant correlation between S. P. CD8+ T cells and Sen. CD8+ T cells in severe & critical COVID-19 patients. (F) Enriched pathways of marker genes and (G) pseudo-time trajectory of TEF CD8+ cells, Sen. CD8+ T cells, Th17 cells, and S. P. CD8+ T cells in severe & critical of COVID-19 patients. (H) Cell-cell communication network and significant ligand-receptor interactions. (I) Diagram of the cell-cell interactions and core transcriptional regulatory relationships of TEF CD8+ cells, Sen. CD8+ T cells, Th17 cells, and S. P. CD8+ T cells.

**Figure 6.**
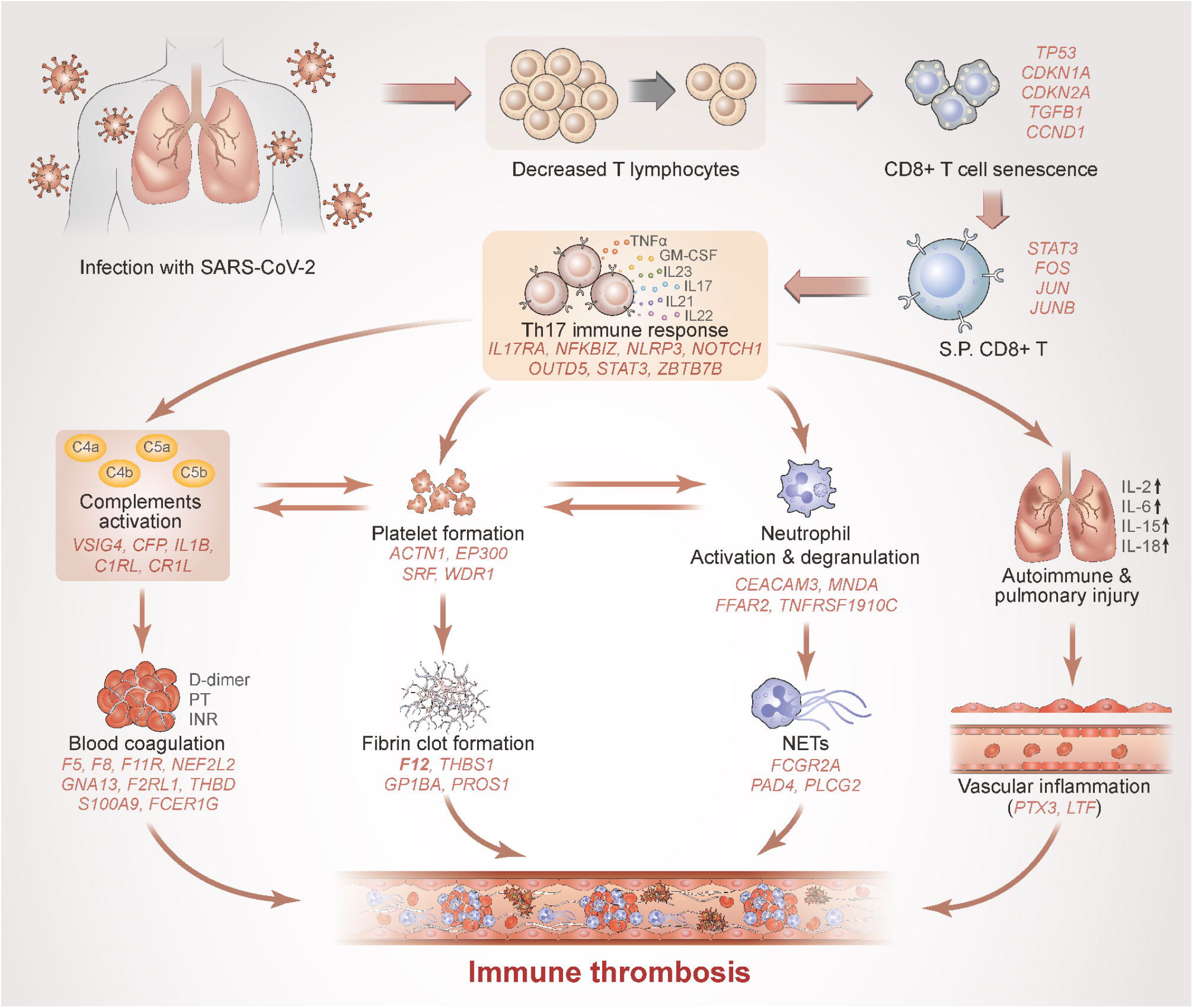
A theoretical view of the S. P. CD8+ T cell contribute to immunothromosis via Th17 response. After the infection with SARS-CoV-2, it causes the decrease of T lymphocytes and the senescence of CD8+ T cells. The senescent CD8+ T cells in severe and critical patients lead to the emergence of a specific phenotype (S. P.) CD8+ T cells, thereby activating the Th17 immune response. Which subsequently causes a series of downstream immunological changes, such as: 1) complement activation and blood hypercoagulation; 2) platelet activation and fibrin clot formation; 3) neutrophil activation, degranulation and NETosis; 4) lung injury and vascular inflammation. All these factors together promote immunothromosis.

## 4. Discussion

SARS-CoV-2 and its variants infection can result in a range of clinical manifestations, from asymptomatic or mild infection to severe COVID-19 that requires hospitalization. T cells have a two-side role in SARS-CoV-2 infection [44]. Activation of T cells promotes resolution of COVID-19 and halts disease progression, with the potential to provide long-term protection against reinfection with SARS-CoV-2. However, in some severe patients, hyperactivation or hypoactivation of T cells, or skewing towards an ineffective differentiation state, may lead to severe disease progression, especially a series of rapid immunological responses such as decreased T lymphocytes, CD8+ T cell senescence, cytokine storm, Th17 immune activation, NETosis, immunothrombosis. Our data showed that these processes were regulated by multiple transcription factors and genes, especially the over-expressed cellular senescence-related genes in severe and critical COVID-19 patients (such as FOS, JUN, NFKBIA, etc.). Infection with the SARS-Cov-2 will cause organ and cellular senescence, and these multiple changes caused by senescence make it hard for the body to effectively defend against virus invasion.

In COVID-19 survivors, persistent post-COVID-19 syndrome (PPCS) presents one or more symptoms: fatigue, dyspnea, memory loss, sleep disorders, and difficulty concentrating. A consistent biological age increases and telomere shortening were found in the post-COVID-19 population [45]. Accumulation of senescent cells increases risk of more severe cases or complications from COVID-19. Immunosenescence causes an increase in baseline levels of multiple cytokines, including IL-1, IL-6, IL-RA, TNF-a, and other senescence-associated secretory phenotype (SASP) factors [46]. Cytokine-mediated senescence and associated inflammation promote and reinforce each another, leading to increased senescence and hyper-inflammation [47]. Animal experiments show that senolytics selectively eliminates virus-induced senescence (VIS) cells, reducing lung infection and inflammation caused by COVID-19 [48].

Our results showed that senescent CD8+ T cells may drive the activation of the S. P. CD8+ T cells in severe and critical COVID-19 patients, further leading to the Th17 cell differentiation and triggering immunothrombosis. We observed the S. P. CD8+ T cells both in PBMC and BAL in severe and critical COVID-19 patients, but not in mild and moderate, convalescence or healthy people. However, the additive effects of SARS-Cov-2 infection and cellular senescence that cause the presence of S. P. CD8+ T cells is unclear. It is noteworthy that whether this process of generating a special class of cells caused by cellular senescence, which in turn drives the downstream immune response, is present in other pandemics.

Dysfunction of CD8+ T cells is the dominant reason for COVID-19 severe progression. Persistent infection and sustained stimulation are factors associated with the exhaustive state in CD8+ T cells [49]. Senescent T cells may damage healthy tissue under severe inflammation caused by COVID-19 [50]. Limiting the expansion or increasing the clearance of senescent CD8+ T cells may be crucial for preventing and ameliorating viral infections, autoimmune disorders and other aging-related diseases [51]. Elderly COVID-19 patients are more likely to accumulate high levels of cellular senescence, leading to uncontrolled SARS-CoV-2-mediated cytokine storms and excessive inflammatory reaction during the early phase of the disease, which in turn promote tissue damage leading to lung failure and multi-tissue dysfunctions, and might negatively impact on vaccine efficacy [52].

As SARS-CoV-2 spreads rapidly around the world, new and comparatively more contagious variants have become a threat to global public health, leading to a surge of COVID-19 in different countries [53]. SARS-CoV-2 Delta variant spreads transplacentally and causes placental cells to exhibit extensive apoptosis, senescence and ferroptosis [54]. The Omicron variant shows abnormal transmission and infectivity and has become the dominant variant in recent months, with partial resistance to therapeutic antibodies and neutralizing activity of convalescent sera [55]. Cellular senescence may be one of the important factors driving the generation of SARS-CoV-2 variants, such as enhancement of RNA-editing enzymes APOBE which resulting in increased mutation frequency [56]. Enhanced anti-aging capacity also contributes to immunity to virus variants. Long-living populations are less susceptible to inflammation and develop more resiliency to COVID-19, may be attributed to the stronger anti-senescence ability [57]. We believed this research further increases our understanding of the mechanisms in COVID-19 variants.

In conclusion, this study revealed that the specific immune response of severe and critical COVID-19 patients due to decreased T lymphocytes and cell senescence may be one of the important causes of immunothrombosis. These findings have an important reference for avoiding the conversion of patients with mild to severe disease. With the continuous mutation of the SARS-CoV-2, the virulence of the variants has weakened but the transmission ability has increased, and the vast majority of infected people are asymptomatic or mild. This study found the presence of S. P. CD8+ cells in severe and critical COVID-19 patients, which might be helpful for quickly identifying those who are likely to develop severe or critical patients from a large population.

## Acknowledgement

This study was supported by the National Natural Science Foundation of China (Nos. 81703166 and 32070676) and the Guangxi Medical University Training Program for Distinguished Young Scholars (to S.Q.A.). The funders had no role in study design, data collection and analysis, decision to publish or preparation of the manuscript.

