## Supplementary material for "Multi-omics integrated analysis reveals a specific phenotype of CD8+ T cell may contribute to immunothromosis via Th17 response in severe and critical COVID-19": Table S1

**Table S1.** Clinical and laboratory characteristics of patients and number of patients included for sequencing and testing.

|  | **COVID-19** | | | **Pneumonia**  **(N=55)** | **Healthy donors**  **(N=36)** |
| --- | --- | --- | --- | --- | --- |
|  | **Mild & Moderate**  **(N=707)** | **Severe**  **(N=448)** | **Critical**  **(N=420)** |  |  |
| ***Clinical & Laboratory characteristics*** | |  |  |  |  |
| Gender, n |  |  |  |  |  |
| Male | 336 (47.5%) | 232 (51.8%) | 271 (64.5%) | 29 (52.7%) | 20 (55.6%) |
| Female | 371 (52.5%) | 216 (48.2%) | 149 (35.5%) | 26 (47.3%) | 16 (44.4%) |
| Age, years | 61 (10-94) | 66 (24-92) | 68 (19-95) | 60 (33-76) | 56 (22-68) |
| White blood cell, ×10^9^/L | 5.6 (4.2-7.3) | 6.3 (4.9-8.7) | 11.1 (7.1-15.6) | 9.0 (6.6-12.7) | 5.1 (4.5-6.9) |
| Neutrophil, ×10^9^/L | 3.7 (2.5-5.4) | 4.1 (2.9-6.3) | 9.8 (5.8-14.5) | 7.0 (5.1-10.2) | 3.5 (2.7-4.1) |
| Lymphocyte, ×10^9^/L | 1.1 (0.8-1.5) | 1.3 (0.9-1.7) | 0.7 (0.4-1.1) | 1.4 (0.9-1.8) | 1.5 (1.3-2.1) |
| Hemoglobin, g/L | 128 (117-141) | 118 (107-130) | 114 (98-129) | 108 (95-122) | 131 (119-139) |
| Platelet, ×10^9^/L | 214 (164-283) | 217 (169-284) | 134 (64-213) | 273 (197-344) | 231 (201-277) |
| Albumin, g/L | 38.0 (34.9-40.8) | 31.6 (28.6-36.5) | 31.1 (26.1-33.7) | 33.1 (30.2-35.7) | 40.1 (37.3-42.7) |
| Globulin, g/L | 27.2 (24.7-29.5) | 29.4 (25.7-38.2) | 34.6 (24.0-41.0) | 34.7 (30.5-38.0) | 29.9 (25.6-32.6) |
| Creatinine, µmol/L | 65.0 (55.0-78.0) | 68.0 (57.5-117.5) | 60.5 (45.3-123.0) | 47.2 (40.5-61.6) | 63 (58-72) |
| Urea nitrogen, mmol/L | 4.3 (3.4-5.4) | 9.1 (5.8-16.8) | 8.4 (5.5-52.4) | 4.6 (3.5-6.2) | 4.8 (3.1-6.3) |
| D-dimer, µg/L | 159 (78-302) | 828 (470-2979) | 2102 (788-7786) | 340 (100-680) | 148 (70-288) |
| Fibrinogen, g/L | 4.0 (3.5-4.6) | 4.3 (3.3-4.8) | 4.3 (3.3-4.8) | 3.8 (3.0-4.6) | 3.4 (2.9-4.6) |
| INR | 1.1 (1.0-1.1) | 1.2 (1.1-1.4) | 1.2 (1.1-1.4) | 1.0 (1.0-1.1) | 1.0 (0.9-1.1) |
| APTT, sec. | 32.3 (29.8-34.8) | 31.2 (26.4-35.0) | 30.2 (25.8-34.9) | 26.5 (23.5-31.4) | 25.4 (23.3-32.5) |
| PT, sec. | 11.8 (11.2-12.4) | 13.0 (11.6-14.0) | 13.2 (11.8-14.8) | 11.8 (11.3-12.4) | 10.9 (10.1-12.8) |
| Outcome, dead | 0 (0%) | 20 (4.5%) | 294 (70.0%) | 0 (0%) | N/A |
| ***Sequencing & Testing*** |  |  |  |  |  |
| Whole blood RNA-seq, n | 30 | 24 | 20 | 18 | 12 |
| Complements, ELISA, n | 30 | 24 | 20 | 18 | 12 |
| Flow cytometry, n | 140 | 32 | 18 | 12 | 6 |

Data are shown as median (IQR) or n/N (%), unless otherwise indicated. APTT, activated partial thromboplastin time; INR, international normalized ratio; PBMC, Peripheral blood mononuclear cell; PT, prothrombin time; RNA-seq, RNA sequencing.
