## Supplementary material for "Multi-omics integrated analysis reveals a specific phenotype of CD8+ T cell may contribute to immunothromosis via Th17 response in severe and critical COVID-19": Table S2

**Table S2**. Up-regulated genes in S. P. CD8+ T cells in PBMC.

| **Gene** | **log2FC** | **Percent in S. P. CD8+ T** | **Percent in other cells** | **P value** | **Adj. P value** |
| --- | --- | --- | --- | --- | --- |
| S100A81 | 2.077 | 0.783 | 0.199 | 0 | 0 |
| S100A91 | 1.790 | 0.672 | 0.214 | 0 | 0 |
| CCL43 | 1.761 | 0.478 | 0.278 | 0 | 0 |
| ZFP364 | 1.753 | 0.830 | 0.613 | 0 | 0 |
| NFKBIA3 | 1.546 | 0.817 | 0.624 | 0 | 0 |
| IGLV3-25 | 1.390 | 0.262 | 0.024 | 0 | 0 |
| FOS4 | 1.385 | 0.834 | 0.501 | 0 | 0 |
| IGHA1 | 1.371 | 0.302 | 0.041 | 0 | 0 |
| IGKC | 1.309 | 0.292 | 0.062 | 0 | 0 |
| PPP1R15A3 | 1.269 | 0.721 | 0.485 | 0 | 0 |
| 44714 | 1.221 | 0.846 | 0.665 | 0 | 0 |
| NR4A23 | 1.196 | 0.467 | 0.273 | 0 | 0 |
| DUSP22 | 1.194 | 0.582 | 0.354 | 0 | 0 |
| KLF66 | 1.140 | 0.865 | 0.700 | 0 | 0 |
| IFI27 | 1.110 | 0.261 | 0.056 | 0 | 0 |
| CD696 | 1.102 | 0.724 | 0.634 | 0 | 0 |
| DUSP13 | 1.089 | 0.822 | 0.646 | 0 | 0 |
| MYADM1 | 1.073 | 0.478 | 0.243 | 0 | 0 |
| LGALS12 | 1.066 | 0.654 | 0.411 | 0 | 0 |
| AC020916.1 | 1.045 | 0.261 | 0.076 | 0 | 0 |
| LYZ | 1.042 | 0.347 | 0.117 | 0 | 0 |
| FOSB3 | 1.031 | 0.447 | 0.233 | 0 | 0 |
| CST3 | 0.988 | 0.329 | 0.108 | 0 | 0 |
| IFI62 | 0.964 | 0.485 | 0.340 | 0 | 0 |
| TNFSF13B | 0.944 | 0.955 | 0.777 | 0 | 0 |
| IFITM31 | 0.924 | 0.452 | 0.291 | 0 | 0 |
| LAG32 | 0.905 | 0.316 | 0.131 | 0 | 0 |
| CST73 | 0.903 | 0.733 | 0.479 | 0 | 0 |
| TSC22D31 | 0.875 | 0.858 | 0.810 | 0 | 0 |
| ADGRE52 | 0.862 | 0.598 | 0.409 | 0 | 0 |
| HSPA51 | 0.861 | 0.633 | 0.497 | 0 | 0 |
| HSP90AA13 | 0.859 | 0.861 | 0.796 | 0 | 0 |
| PPBP | 0.859 | 0.291 | 0.098 | 0 | 0 |
| AD000090.11 | 0.851 | 0.961 | 0.789 | 0 | 0 |
| CSRNP12 | 0.831 | 0.343 | 0.178 | 0 | 0 |
| CD83 | 0.825 | 0.254 | 0.094 | 0 | 0 |
| HLA-DPA13 | 0.817 | 0.563 | 0.390 | 0 | 0 |
| PLEK3 | 0.812 | 0.522 | 0.309 | 0 | 0 |
| CD743 | 0.767 | 0.895 | 0.771 | 0 | 0 |
| GZMB2 | 0.754 | 0.575 | 0.340 | 0 | 0 |
| LINC024461 | 0.737 | 0.292 | 0.135 | 0 | 0 |
| HLA-DPB13 | 0.730 | 0.540 | 0.380 | 0 | 0 |
| IRF11 | 0.726 | 0.621 | 0.461 | 0 | 0 |
| IFITM12 | 0.715 | 0.951 | 0.947 | 0 | 0 |
| JUND1 | 0.713 | 0.501 | 0.290 | 0 | 0 |
| HLA-DRA3 | 0.712 | 0.331 | 0.174 | 0 | 0 |
| KLRD11 | 0.708 | 0.548 | 0.336 | 0 | 0 |
| CCL53 | 0.701 | 0.809 | 0.520 | 0 | 0 |
| PRF12 | 0.694 | 0.597 | 0.400 | 0 | 0 |
| NKG73 | 0.691 | 0.777 | 0.496 | 0 | 0 |
| GZMH3 | 0.678 | 0.642 | 0.380 | 0 | 0 |
| SOCS1 | 0.669 | 0.250 | 0.119 | 0 | 0 |
| DAAM1 | 0.667 | 0.612 | 0.471 | 0 | 0 |
| SRGN3 | 0.667 | 0.856 | 0.692 | 0 | 0 |
| KLRG13 | 0.664 | 0.423 | 0.256 | 0 | 0 |
| CYBA1 | 0.631 | 0.815 | 0.738 | 0 | 0 |
| ZFP36L24 | 0.586 | 0.869 | 0.803 | 0 | 0 |
| S100A105 | 0.562 | 0.834 | 0.762 | 0 | 0 |
| FTH16 | 0.561 | 0.982 | 0.954 | 0 | 0 |
| CTSW2 | 0.528 | 0.725 | 0.523 | 0 | 0 |
| UBB1 | 0.514 | 0.821 | 0.774 | 0 | 0 |
| ZNF688 | 0.513 | 0.782 | 0.683 | 0 | 0 |
| TAGLN22 | 0.511 | 0.809 | 0.738 | 0 | 0 |
| S100A63 | 0.494 | 0.911 | 0.887 | 0 | 0 |
| CALM13 | 0.489 | 0.942 | 0.924 | 0 | 0 |
| CLDN111 | 0.476 | 0.803 | 0.648 | 0 | 0 |
| PFN13 | 0.472 | 0.967 | 0.954 | 0 | 0 |
| GAPDH3 | 0.455 | 0.924 | 0.908 | 0 | 0 |
| H3F3B | 0.437 | 0.963 | 0.954 | 0 | 0 |
| GNLY2 | 0.421 | 0.629 | 0.381 | 0 | 0 |
| RNF2132 | 0.658 | 0.685 | 0.574 | 4.00E-307 | 1.10E-302 |
| CLIC13 | 0.569 | 0.747 | 0.634 | 1.50E-295 | 4.00E-291 |
| XAF11 | 0.832 | 0.448 | 0.304 | 3.50E-295 | 9.30E-291 |
| SH3BGRL33 | 0.408 | 0.952 | 0.929 | 1.00E-294 | 2.80E-290 |
| GZMA3 | 0.476 | 0.659 | 0.444 | 5.30E-285 | 1.40E-280 |
| TYROBP1 | 0.724 | 0.419 | 0.248 | 1.30E-282 | 3.40E-278 |
| ISG152 | 0.935 | 0.484 | 0.352 | 2.70E-275 | 7.30E-271 |
| UBC2 | 0.372 | 0.929 | 0.919 | 5.60E-275 | 1.50E-270 |
| ISG201 | 0.698 | 0.648 | 0.560 | 3.50E-268 | 9.30E-264 |
| LRRFIP12 | 0.586 | 0.630 | 0.524 | 9.70E-266 | 2.60E-261 |
| ID23 | 0.647 | 0.492 | 0.340 | 1.60E-264 | 4.20E-260 |
| NEAT1 | 0.801 | 0.626 | 0.502 | 4.60E-264 | 1.20E-259 |
| REL1 | 0.667 | 0.452 | 0.314 | 2.00E-259 | 5.40E-255 |
| TNFAIP35 | 1.162 | 0.580 | 0.485 | 1.40E-255 | 3.60E-251 |
| DNAJA1 | 0.645 | 0.485 | 0.358 | 1.70E-250 | 4.60E-246 |
| YPEL52 | 0.763 | 0.502 | 0.383 | 3.00E-249 | 7.80E-245 |
| EMP32 | 0.535 | 0.818 | 0.791 | 1.20E-239 | 3.30E-235 |
| AL024508.2 | 0.575 | 0.293 | 0.168 | 4.70E-238 | 1.20E-233 |
| AC092171.3 | 0.571 | 0.260 | 0.142 | 2.40E-237 | 6.30E-233 |
| CD8A3 | 0.478 | 0.462 | 0.290 | 1.30E-234 | 3.40E-230 |
| TNRC6B | 0.534 | 0.616 | 0.510 | 1.30E-229 | 3.60E-225 |
| HCST3 | 0.435 | 0.851 | 0.803 | 5.80E-224 | 1.50E-219 |
| SERF21 | 0.345 | 0.939 | 0.949 | 3.50E-223 | 9.30E-219 |
| S100A115 | 0.634 | 0.592 | 0.479 | 1.50E-222 | 3.90E-218 |
| AMFR | 0.421 | 0.771 | 0.684 | 6.30E-222 | 1.70E-217 |
| HLA-DRB53 | 0.745 | 0.428 | 0.290 | 1.40E-220 | 3.70E-216 |
| DDIT3 | 0.584 | 0.259 | 0.144 | 5.40E-218 | 1.40E-213 |
| HSP90B11 | 0.581 | 0.646 | 0.562 | 5.10E-215 | 1.40E-210 |
| BTG22 | 0.816 | 0.476 | 0.365 | 6.10E-208 | 1.60E-203 |
| AC106820.2 | 0.593 | 0.411 | 0.289 | 2.90E-205 | 7.70E-201 |
| SERTAD1 | 0.549 | 0.275 | 0.158 | 6.60E-205 | 1.80E-200 |
| LY6E2 | 0.609 | 0.748 | 0.707 | 4.20E-204 | 1.10E-199 |
| ZEB23 | 0.545 | 0.395 | 0.256 | 8.50E-203 | 2.30E-198 |
| GADD45B2 | 0.850 | 0.502 | 0.386 | 2.90E-202 | 7.70E-198 |
| ARID5B | 0.350 | 0.882 | 0.829 | 3.20E-202 | 8.50E-198 |
| S100A45 | 0.386 | 0.911 | 0.834 | 3.10E-201 | 8.10E-197 |
| JUNB6 | 0.681 | 0.799 | 0.755 | 1.90E-200 | 4.90E-196 |
| PSMB93 | 0.552 | 0.687 | 0.632 | 1.30E-199 | 3.50E-195 |
| HSP90AB11 | 0.465 | 0.785 | 0.772 | 1.10E-192 | 3.00E-188 |
| HLA-E2 | 0.295 | 0.963 | 0.970 | 8.40E-192 | 2.20E-187 |
| TYMP | 0.561 | 0.255 | 0.148 | 1.90E-190 | 5.00E-186 |
| GSTP12 | 0.510 | 0.608 | 0.511 | 2.40E-186 | 6.30E-182 |
| SH2D2A1 | 0.549 | 0.349 | 0.233 | 2.90E-185 | 7.60E-181 |
| CYCS | 0.574 | 0.512 | 0.420 | 4.20E-184 | 1.10E-179 |
| SON | 0.402 | 0.755 | 0.711 | 9.00E-180 | 2.40E-175 |
| CTSC3 | 0.553 | 0.502 | 0.396 | 3.80E-177 | 1.00E-172 |
| IL323 | 0.346 | 0.922 | 0.906 | 6.20E-175 | 1.70E-170 |
| BRD21 | 0.564 | 0.514 | 0.415 | 1.10E-170 | 2.90E-166 |
| IRF71 | 0.614 | 0.336 | 0.228 | 1.00E-168 | 2.70E-164 |
| RELB | 0.478 | 0.254 | 0.151 | 3.60E-165 | 9.60E-161 |
| MCL1 | 0.565 | 0.501 | 0.408 | 1.80E-164 | 4.80E-160 |
| PNRC1 | 0.397 | 0.776 | 0.752 | 3.20E-164 | 8.50E-160 |
| NFKBIZ1 | 0.666 | 0.390 | 0.283 | 3.90E-164 | 1.00E-159 |
| BHLHE404 | 0.578 | 0.292 | 0.185 | 1.40E-161 | 3.80E-157 |
| HLA-DRB13 | 0.455 | 0.311 | 0.202 | 3.40E-153 | 9.10E-149 |
| SQSTM11 | 0.582 | 0.484 | 0.402 | 1.80E-149 | 4.70E-145 |
| PPIB3 | 0.432 | 0.738 | 0.713 | 2.00E-149 | 5.20E-145 |
| EIF5 | 0.497 | 0.506 | 0.420 | 6.30E-149 | 1.70E-144 |
| FGR2 | 0.469 | 0.268 | 0.168 | 1.20E-146 | 3.20E-142 |
| AKAP131 | 0.485 | 0.507 | 0.420 | 3.50E-146 | 9.30E-142 |
| ACTB2 | 0.279 | 0.998 | 0.995 | 1.80E-143 | 4.80E-139 |
| ANXA16 | 0.408 | 0.703 | 0.619 | 5.70E-143 | 1.50E-138 |
| KLF2 | 0.461 | 0.600 | 0.525 | 2.60E-141 | 6.90E-137 |
| MYL63 | 0.338 | 0.872 | 0.875 | 3.00E-140 | 7.80E-136 |
| HIST1H4C | 0.485 | 0.527 | 0.452 | 3.70E-140 | 9.70E-136 |
| CDK2AP22 | 0.551 | 0.373 | 0.278 | 3.70E-139 | 9.80E-135 |
| DDX24 | 0.433 | 0.587 | 0.523 | 5.80E-138 | 1.50E-133 |
| MYL12A2 | 0.320 | 0.925 | 0.937 | 8.90E-137 | 2.30E-132 |
| HERPUD11 | 0.517 | 0.363 | 0.266 | 8.60E-135 | 2.30E-130 |
| CD993 | 0.351 | 0.758 | 0.710 | 1.40E-134 | 3.80E-130 |
| CD533 | 0.457 | 0.601 | 0.543 | 3.40E-134 | 9.00E-130 |
| FCGR3A1 | 0.346 | 0.356 | 0.235 | 5.80E-134 | 1.50E-129 |
| MTPN1 | 0.479 | 0.386 | 0.292 | 2.70E-133 | 7.20E-129 |
| STK17A | 0.443 | 0.512 | 0.429 | 1.70E-132 | 4.60E-128 |
| GZMM3 | 0.402 | 0.638 | 0.565 | 3.00E-131 | 7.90E-127 |
| PSME21 | 0.468 | 0.632 | 0.589 | 1.30E-129 | 3.50E-125 |
| TENT5C2 | 0.481 | 0.273 | 0.180 | 2.30E-126 | 6.20E-122 |
| MX1 | 0.540 | 0.312 | 0.220 | 3.80E-126 | 1.00E-121 |
| TCF251 | 0.428 | 0.551 | 0.476 | 2.30E-125 | 6.00E-121 |
| B4GALT11 | 0.447 | 0.465 | 0.381 | 4.20E-125 | 1.10E-120 |
| FCRL62 | 0.403 | 0.295 | 0.196 | 1.10E-123 | 2.80E-119 |
| CCDC85B1 | 0.436 | 0.517 | 0.442 | 7.30E-123 | 1.90E-118 |
| BST21 | 0.622 | 0.456 | 0.389 | 3.80E-122 | 1.00E-117 |
| HIST1H1C | 0.421 | 0.258 | 0.168 | 2.00E-121 | 5.40E-117 |
| SP110 | 0.489 | 0.432 | 0.349 | 1.90E-118 | 5.00E-114 |
| RAB292 | 0.462 | 0.346 | 0.259 | 1.00E-117 | 2.80E-113 |
| AGTRAP1 | 0.452 | 0.272 | 0.188 | 4.10E-115 | 1.10E-110 |
| TUBA1A | 0.535 | 0.475 | 0.406 | 2.40E-114 | 6.30E-110 |
| CD8B4 | 0.276 | 0.360 | 0.243 | 2.90E-114 | 7.80E-110 |
| ARPC5L3 | 0.444 | 0.427 | 0.344 | 2.10E-112 | 5.50E-108 |
| SYNE13 | 0.493 | 0.435 | 0.352 | 2.30E-112 | 6.10E-108 |
| MT2A3 | 0.383 | 0.496 | 0.422 | 1.10E-111 | 2.80E-107 |
| C12orf753 | 0.385 | 0.433 | 0.335 | 4.70E-111 | 1.30E-106 |
| ATP6V0C | 0.391 | 0.524 | 0.457 | 3.50E-108 | 9.40E-104 |
| CMC12 | 0.313 | 0.340 | 0.239 | 1.00E-107 | 2.70E-103 |
| DDIT43 | 0.574 | 0.443 | 0.361 | 1.70E-106 | 4.50E-102 |
| CISD32 | 0.452 | 0.255 | 0.174 | 4.70E-106 | 1.20E-101 |
| ABI32 | 0.426 | 0.254 | 0.171 | 6.50E-106 | 1.70E-101 |
| RGCC6 | 0.801 | 0.312 | 0.234 | 1.20E-103 | 3.30E-99 |
| CX3CR12 | 0.346 | 0.280 | 0.188 | 3.80E-102 | 1.01E-97 |
| SAT12 | 0.458 | 0.540 | 0.481 | 8.50E-102 | 2.26E-97 |
| SYTL32 | 0.429 | 0.312 | 0.229 | 4.30E-100 | 1.15E-95 |
| KLRF11 | 0.328 | 0.250 | 0.163 | 8.80E-100 | 2.34E-95 |
| ALOX5AP1 | 0.534 | 0.430 | 0.356 | 1.30E-99 | 3.37E-95 |
| DNAJC13 | 0.430 | 0.272 | 0.193 | 2.79E-98 | 7.40E-94 |
| FLNA3 | 0.347 | 0.596 | 0.521 | 9.49E-98 | 2.52E-93 |
| RUNX33 | 0.419 | 0.353 | 0.271 | 6.74E-97 | 1.79E-92 |
| VIM4 | 0.286 | 0.811 | 0.788 | 9.68E-96 | 2.57E-91 |
| MYO1G2 | 0.415 | 0.411 | 0.336 | 8.41E-94 | 2.23E-89 |
| DNAJB13 | 0.753 | 0.531 | 0.496 | 1.58E-93 | 4.19E-89 |
| AHNAK5 | 0.293 | 0.619 | 0.523 | 2.69E-93 | 7.13E-89 |
| TTC381 | 0.349 | 0.252 | 0.172 | 5.77E-92 | 1.53E-87 |
| IQGAP12 | 0.377 | 0.460 | 0.386 | 4.35E-89 | 1.15E-84 |
| ZNF3312 | 0.587 | 0.300 | 0.224 | 6.89E-89 | 1.83E-84 |
| MTRNR2L6 | 0.493 | 0.380 | 0.311 | 6.85E-88 | 1.82E-83 |
| EIF5A | 0.350 | 0.586 | 0.549 | 2.64E-86 | 7.01E-82 |
| TUBB4B1 | 0.365 | 0.311 | 0.234 | 3.54E-85 | 9.38E-81 |
| MYO1F3 | 0.412 | 0.354 | 0.276 | 1.00E-84 | 2.66E-80 |
| CDC42EP33 | 0.400 | 0.269 | 0.196 | 1.78E-84 | 4.73E-80 |
| EIF2AK2 | 0.496 | 0.317 | 0.249 | 2.35E-84 | 6.23E-80 |
| GNB21 | 0.374 | 0.438 | 0.371 | 8.01E-84 | 2.12E-79 |
| BTG13 | 0.392 | 0.862 | 0.876 | 5.72E-83 | 1.52E-78 |
| IKZF33 | 0.424 | 0.334 | 0.262 | 1.03E-82 | 2.72E-78 |
| TMBIM61 | 0.346 | 0.643 | 0.630 | 4.23E-82 | 1.12E-77 |
| GNG51 | 0.427 | 0.502 | 0.460 | 4.44E-82 | 1.18E-77 |
| SP1001 | 0.387 | 0.525 | 0.480 | 1.35E-81 | 3.57E-77 |
| CD631 | 0.389 | 0.374 | 0.301 | 3.12E-81 | 8.27E-77 |
| IDS1 | 0.393 | 0.420 | 0.354 | 3.40E-80 | 9.02E-76 |
| RBM8A | 0.368 | 0.491 | 0.442 | 4.32E-79 | 1.14E-74 |
| TUBA1B1 | 0.371 | 0.479 | 0.421 | 1.13E-78 | 3.00E-74 |
| EFHD22 | 0.306 | 0.344 | 0.262 | 2.99E-78 | 7.92E-74 |
| PDIA31 | 0.363 | 0.583 | 0.550 | 3.65E-78 | 9.69E-74 |
| STAT3 | 0.397 | 0.435 | 0.377 | 2.40E-75 | 6.37E-71 |
| RSBN1L | 0.415 | 0.318 | 0.254 | 2.59E-75 | 6.88E-71 |
| RAC22 | 0.254 | 0.832 | 0.840 | 6.68E-75 | 1.77E-70 |
| RASSF12 | 0.362 | 0.332 | 0.261 | 1.54E-74 | 4.08E-70 |
| GNG22 | 0.333 | 0.476 | 0.412 | 1.97E-73 | 5.21E-69 |
| TPST21 | 0.362 | 0.335 | 0.264 | 6.43E-73 | 1.71E-68 |
| SMAD2 | 0.395 | 0.400 | 0.342 | 1.11E-72 | 2.95E-68 |
| DRAP11 | 0.363 | 0.538 | 0.499 | 1.43E-70 | 3.80E-66 |
| TPM32 | 0.284 | 0.750 | 0.762 | 4.56E-69 | 1.21E-64 |
| IFI44L | 0.379 | 0.269 | 0.202 | 1.47E-66 | 3.89E-62 |
| PLAC82 | 0.397 | 0.445 | 0.395 | 3.29E-65 | 8.74E-61 |
| CCND31 | 0.287 | 0.668 | 0.664 | 4.43E-65 | 1.17E-60 |
| NF1 | 0.366 | 0.339 | 0.278 | 6.19E-65 | 1.64E-60 |
| TUBA4A2 | 0.436 | 0.394 | 0.343 | 1.11E-64 | 2.94E-60 |
| ACTN43 | 0.359 | 0.282 | 0.219 | 3.82E-63 | 1.01E-58 |
| ARL4C2 | 0.329 | 0.514 | 0.472 | 4.73E-63 | 1.25E-58 |
| IER2 | 0.358 | 0.587 | 0.543 | 5.13E-62 | 1.36E-57 |
| TPM42 | 0.373 | 0.291 | 0.230 | 6.79E-62 | 1.80E-57 |
| SELENOK | 0.339 | 0.429 | 0.380 | 9.12E-62 | 2.42E-57 |
| APOBEC3G3 | 0.342 | 0.337 | 0.275 | 5.55E-61 | 1.47E-56 |
| PPP1R183 | 0.342 | 0.455 | 0.407 | 2.49E-60 | 6.61E-56 |
| MACF11 | 0.373 | 0.474 | 0.427 | 7.88E-60 | 2.09E-55 |
| CHCHD101 | 0.341 | 0.325 | 0.267 | 1.85E-59 | 4.90E-55 |
| ANXA62 | 0.362 | 0.548 | 0.528 | 8.57E-59 | 2.27E-54 |
| SLC2A36 | 0.448 | 0.433 | 0.381 | 1.07E-58 | 2.83E-54 |
| SHISA51 | 0.363 | 0.474 | 0.440 | 3.43E-58 | 9.11E-54 |
| TLN12 | 0.353 | 0.350 | 0.292 | 3.76E-58 | 9.98E-54 |
| XBP11 | 0.325 | 0.421 | 0.368 | 6.63E-58 | 1.76E-53 |
| S1PR52 | 0.279 | 0.261 | 0.195 | 7.36E-58 | 1.95E-53 |
| GUK12 | 0.292 | 0.586 | 0.565 | 7.32E-57 | 1.94E-52 |
| CDC42SE11 | 0.326 | 0.462 | 0.419 | 1.06E-56 | 2.81E-52 |
| RARRES33 | 0.348 | 0.642 | 0.648 | 1.13E-56 | 3.00E-52 |
| YWHAB1 | 0.270 | 0.734 | 0.750 | 3.35E-56 | 8.89E-52 |
| ELF1 | 0.325 | 0.473 | 0.438 | 5.99E-56 | 1.59E-51 |
| EZR4 | 0.391 | 0.453 | 0.416 | 7.47E-56 | 1.98E-51 |
| IFRD1 | 0.345 | 0.251 | 0.193 | 4.28E-55 | 1.14E-50 |
| UTRN1 | 0.385 | 0.374 | 0.324 | 2.63E-54 | 6.96E-50 |
| PRRC2C1 | 0.295 | 0.614 | 0.600 | 6.19E-54 | 1.64E-49 |
| PRELID11 | 0.315 | 0.431 | 0.383 | 7.66E-54 | 2.03E-49 |
| WAS1 | 0.325 | 0.419 | 0.375 | 2.51E-53 | 6.67E-49 |
| SAMD33 | 0.321 | 0.347 | 0.288 | 2.96E-53 | 7.85E-49 |
| ADAR | 0.350 | 0.410 | 0.364 | 1.14E-52 | 3.01E-48 |
| DOK22 | 0.342 | 0.384 | 0.334 | 1.56E-52 | 4.13E-48 |
| RHOH2 | 0.437 | 0.465 | 0.446 | 1.39E-50 | 3.69E-46 |
| ZBTB383 | 0.351 | 0.287 | 0.234 | 1.52E-49 | 4.03E-45 |
| CFLAR1 | 0.322 | 0.448 | 0.408 | 2.04E-49 | 5.42E-45 |
| TOB13 | 0.459 | 0.315 | 0.268 | 3.46E-49 | 9.19E-45 |
| CTSS1 | 0.382 | 0.347 | 0.305 | 3.70E-49 | 9.80E-45 |
| PSMA71 | 0.315 | 0.571 | 0.569 | 4.03E-49 | 1.07E-44 |
| FAM177A1 | 0.334 | 0.279 | 0.225 | 6.21E-49 | 1.65E-44 |
| SRRM2 | 0.316 | 0.455 | 0.413 | 6.53E-49 | 1.73E-44 |
| NFATC22 | 0.323 | 0.289 | 0.234 | 1.53E-48 | 4.05E-44 |
| PTPN61 | 0.362 | 0.386 | 0.345 | 6.51E-48 | 1.73E-43 |
| PTPN72 | 0.335 | 0.264 | 0.213 | 1.33E-47 | 3.51E-43 |
| DIAPH12 | 0.322 | 0.292 | 0.240 | 1.81E-47 | 4.81E-43 |
| SMG1 | 0.323 | 0.508 | 0.479 | 4.09E-47 | 1.08E-42 |
| STK102 | 0.338 | 0.343 | 0.296 | 1.70E-46 | 4.52E-42 |
| MTDH | 0.316 | 0.467 | 0.440 | 5.29E-46 | 1.40E-41 |
| CALR2 | 0.324 | 0.594 | 0.588 | 3.06E-45 | 8.13E-41 |
| KMT2E | 0.283 | 0.553 | 0.539 | 3.84E-45 | 1.02E-40 |
| NCOR1 | 0.329 | 0.417 | 0.385 | 2.56E-42 | 6.78E-38 |
| G3BP2 | 0.355 | 0.398 | 0.364 | 4.75E-42 | 1.26E-37 |
| TIMP13 | 0.338 | 0.278 | 0.231 | 1.22E-41 | 3.23E-37 |
| SNX6 | 0.324 | 0.374 | 0.339 | 2.17E-41 | 5.74E-37 |
| PRKCB1 | 0.325 | 0.258 | 0.212 | 3.13E-41 | 8.31E-37 |
| RTN4 | 0.320 | 0.343 | 0.305 | 1.16E-40 | 3.08E-36 |
| RSRP1 | 0.342 | 0.545 | 0.541 | 1.36E-40 | 3.62E-36 |
| APBB1IP1 | 0.316 | 0.416 | 0.385 | 2.00E-40 | 5.30E-36 |
| TMEM50A2 | 0.288 | 0.485 | 0.466 | 2.70E-40 | 7.15E-36 |
| CCND2 | 0.336 | 0.251 | 0.207 | 7.20E-40 | 1.91E-35 |
| KRTCAP22 | 0.271 | 0.581 | 0.585 | 3.71E-39 | 9.83E-35 |
| ARHGAP301 | 0.325 | 0.372 | 0.336 | 4.68E-39 | 1.24E-34 |
| CHST123 | 0.326 | 0.297 | 0.253 | 1.50E-38 | 3.98E-34 |
| CAST2 | 0.284 | 0.475 | 0.446 | 8.52E-38 | 2.26E-33 |
| JPT13 | 0.311 | 0.276 | 0.234 | 1.04E-37 | 2.75E-33 |
| CARD161 | 0.334 | 0.367 | 0.334 | 1.98E-37 | 5.26E-33 |
| SH3BP11 | 0.297 | 0.250 | 0.207 | 3.74E-37 | 9.91E-33 |
| SRSF71 | 0.274 | 0.605 | 0.610 | 4.59E-37 | 1.22E-32 |
| VASP2 | 0.309 | 0.314 | 0.275 | 6.03E-37 | 1.60E-32 |
| KRAS | 0.308 | 0.269 | 0.226 | 6.92E-37 | 1.83E-32 |
| PRR131 | 0.260 | 0.620 | 0.634 | 1.08E-36 | 2.87E-32 |
| PHF20 | 0.299 | 0.324 | 0.285 | 2.98E-36 | 7.91E-32 |
| DNMT3A | 0.319 | 0.297 | 0.258 | 1.18E-35 | 3.14E-31 |
| ATP6V0E11 | 0.286 | 0.544 | 0.544 | 1.70E-35 | 4.51E-31 |
| NAPA1 | 0.307 | 0.344 | 0.311 | 5.69E-35 | 1.51E-30 |
| ROCK11 | 0.305 | 0.341 | 0.304 | 6.20E-35 | 1.64E-30 |
| SYNE23 | 0.332 | 0.476 | 0.450 | 2.56E-34 | 6.78E-30 |
| ITGAL3 | 0.284 | 0.336 | 0.295 | 4.30E-34 | 1.14E-29 |
| SLFN5 | 0.355 | 0.327 | 0.293 | 1.92E-33 | 5.09E-29 |
| MBP1 | 0.278 | 0.446 | 0.421 | 1.93E-33 | 5.12E-29 |
| BCAS2 | 0.298 | 0.327 | 0.290 | 2.78E-33 | 7.38E-29 |
| IFI161 | 0.321 | 0.327 | 0.292 | 2.93E-33 | 7.77E-29 |
| PSAP2 | 0.276 | 0.473 | 0.455 | 4.01E-33 | 1.06E-28 |
| KTN1 | 0.300 | 0.353 | 0.322 | 5.93E-33 | 1.57E-28 |
| ZFP36L1 | 0.356 | 0.469 | 0.453 | 2.71E-32 | 7.18E-28 |
| SEC61B2 | 0.264 | 0.504 | 0.497 | 8.79E-32 | 2.33E-27 |
| TXN2 | 0.292 | 0.371 | 0.344 | 1.50E-31 | 3.98E-27 |
| FAM96B1 | 0.287 | 0.361 | 0.334 | 2.87E-31 | 7.61E-27 |
| FAM126B | 0.287 | 0.315 | 0.282 | 6.53E-31 | 1.73E-26 |
| GNAI21 | 0.289 | 0.324 | 0.290 | 8.29E-31 | 2.20E-26 |
| CD552 | 0.397 | 0.301 | 0.268 | 1.22E-30 | 3.23E-26 |
| GNAS | 0.276 | 0.399 | 0.372 | 2.33E-30 | 6.17E-26 |
| REEP53 | 0.285 | 0.305 | 0.273 | 5.78E-29 | 1.53E-24 |
| TGFB13 | 0.270 | 0.365 | 0.336 | 1.46E-28 | 3.88E-24 |
| SELENOW1 | 0.268 | 0.507 | 0.507 | 1.55E-28 | 4.11E-24 |
| VAMP51 | 0.291 | 0.288 | 0.255 | 1.76E-28 | 4.67E-24 |
| RBM6 | 0.295 | 0.297 | 0.263 | 1.86E-28 | 4.92E-24 |
| RSRC2 | 0.276 | 0.352 | 0.321 | 2.45E-28 | 6.49E-24 |
| UBE2L61 | 0.347 | 0.375 | 0.358 | 6.06E-28 | 1.61E-23 |
| UHMK11 | 0.270 | 0.273 | 0.238 | 7.55E-28 | 2.00E-23 |
| PSMB102 | 0.271 | 0.474 | 0.465 | 1.07E-27 | 2.83E-23 |
| JMJD1C | 0.291 | 0.282 | 0.246 | 2.80E-27 | 7.41E-23 |
| NSD31 | 0.256 | 0.399 | 0.375 | 3.94E-27 | 1.05E-22 |
| MYDGF1 | 0.279 | 0.275 | 0.244 | 1.43E-26 | 3.80E-22 |
| GLIPR22 | 0.265 | 0.257 | 0.223 | 2.55E-26 | 6.77E-22 |
| PPP1R12A1 | 0.268 | 0.345 | 0.319 | 4.57E-26 | 1.21E-21 |
| ITSN2 | 0.258 | 0.303 | 0.270 | 6.50E-26 | 1.72E-21 |
| RHOG1 | 0.278 | 0.418 | 0.406 | 1.41E-25 | 3.75E-21 |
| TAF1D | 0.269 | 0.389 | 0.370 | 2.11E-25 | 5.59E-21 |
| DDX21 | 0.250 | 0.355 | 0.329 | 2.45E-25 | 6.50E-21 |
| ZNF24 | 0.268 | 0.263 | 0.230 | 3.07E-25 | 8.15E-21 |
| RSF1 | 0.272 | 0.324 | 0.297 | 4.12E-25 | 1.09E-20 |
| NUCKS1 | 0.267 | 0.448 | 0.439 | 1.19E-24 | 3.15E-20 |
| HERPUD2 | 0.266 | 0.267 | 0.235 | 1.57E-24 | 4.15E-20 |
| PNN | 0.259 | 0.376 | 0.354 | 1.63E-24 | 4.32E-20 |
| AKAP9 | 0.291 | 0.300 | 0.272 | 5.55E-24 | 1.47E-19 |
| GTF2B | 0.264 | 0.262 | 0.229 | 5.63E-24 | 1.49E-19 |
| RPL22L1 | 0.277 | 0.397 | 0.382 | 1.05E-23 | 2.79E-19 |
| AC016831.7 | 0.287 | 0.322 | 0.296 | 3.14E-23 | 8.32E-19 |
| DEK1 | 0.259 | 0.382 | 0.366 | 1.27E-21 | 3.38E-17 |
| CASP13 | 0.276 | 0.257 | 0.230 | 2.23E-21 | 5.90E-17 |
| TRIM22 | 0.263 | 0.380 | 0.363 | 8.08E-21 | 2.14E-16 |
| GBP53 | 0.269 | 0.288 | 0.259 | 8.53E-21 | 2.26E-16 |
| RPN11 | 0.255 | 0.330 | 0.310 | 8.96E-21 | 2.38E-16 |
| RNF149 | 0.263 | 0.332 | 0.312 | 9.44E-21 | 2.50E-16 |
| CHD1 | 0.280 | 0.271 | 0.242 | 2.10E-20 | 5.56E-16 |
| PDIA61 | 0.264 | 0.281 | 0.257 | 2.61E-20 | 6.91E-16 |
| KDM2A | 0.253 | 0.250 | 0.222 | 4.37E-20 | 1.16E-15 |
| UQCRQ1 | 0.256 | 0.416 | 0.412 | 5.48E-20 | 1.45E-15 |
| RASAL32 | 0.250 | 0.299 | 0.275 | 9.06E-20 | 2.40E-15 |
| NDUFAF3 | 0.265 | 0.277 | 0.255 | 4.13E-19 | 1.10E-14 |
| DNAJB14 | 0.255 | 0.289 | 0.266 | 3.29E-18 | 8.74E-14 |
| HSPE1 | 0.285 | 0.375 | 0.370 | 8.29E-17 | 2.20E-12 |
| APOL61 | 0.264 | 0.258 | 0.238 | 4.34E-16 | 1.15E-11 |
| ASH1L | 0.264 | 0.254 | 0.235 | 5.98E-13 | 1.59E-08 |
